# The hippocampal film-editor: sensitivity and specificity to event boundaries in continuous experience

**DOI:** 10.1101/273409

**Authors:** Aya Ben-Yakov, Richard Henson

**Affiliations:** MRC Cognition and Brain Sciences Unit, University of Cambridge, CB2 7EF

## Abstract

The function of the human hippocampus is normally investigated by experimental manipulation of discrete events. Less is unknown about what triggers hippocampal activity during more naturalistic, continuous experience. We hypothesized that the hippocampus would be sensitive to the occurrence of event boundaries, i.e. moments in time identified by observers as a transition between events. To address this, we analysed functional MRI data from two groups: one (N=253, 131 female) who viewed an 8.5min film and another (N=15, 6 female) who viewed a 120min film. We observed a strong hippocampal response at boundaries defined by independent observers, which was modulated by boundary strength (the number of observers that identified each boundary). In the longer film, there were sufficient boundaries to show that this modulation remained after covarying out a large number of perceptual factors. The hippocampus was the only brain region whose response showed a significant monotonic increase with boundary strength in both films, suggesting that modulation by boundary strength is selective to the hippocampus. This hypothesis-driven approach was complemented by a data-driven approach, in which we identified hippocampal-events as moments in time with the strongest hippocampal activity: The correspondence between these hippocampal-events and event boundaries was highly-significant, revealing that the hippocampal response is not only sensitive, but also specific to event boundaries. We conclude that the hippocampus plays an important role in segmenting the continuous experience that is typical of naturalistic settings.

**Significance statement:** Recent years have seen the field of human neuroscience research transitioning from experiments with simple stimuli to the study of more complex and naturalistic experience. Nonetheless, our understanding of the function of many brain regions, such as the hippocampus, is based primarily on the study of brief, discrete events. As a result, we know little of what triggers hippocampal activity in real-life settings, when we are exposed to a continuous stream of information. When does the hippocampus “decide” to respond during the encoding of naturalistic experience? We reveal here that hippocampal activity measured by fMRI during film-watching is both sensitive and specific to event boundaries, identifying a potential mechanism whereby event boundaries shape experience by modulation of hippocampal activity.

## Introduction

The hippocampus is, perhaps, one of the most widely-studied regions in the human brain, and research has suggested that it has many roles. The role revealed by each study depends on the specific experimental design and comparison of interest. For example, one set of studies may find the hippocampus responds more strongly to subsequently-remembered than subsequently-forgotten events (Wagner et al., 1998; Kim, 2011), while others find it responds more strongly to novel events than previously-encountered events (Tulving et al., 1996; Kumaran and Maguire, 2009). The strength of these experiments lies in their ability to probe a specific dimension, comparing the hippocampal response to events in two conditions. Yet real life provides us with a continuous stream of complex information – what is an “event” in this context? In other words, what triggers hippocampal activity in naturalistic settings, when we do not present the hippocampus with discretised events?

According to Event Segmentation Theory (Zacks et al., 2007; Kurby and Zacks, 2008), we naturally segment continuous experience into events, and this segmentation is driven by moments in time when prediction of the immediate future fails (event boundaries). Segmentation affects not only our perception of the experience, but its subsequent organisation in long-term memory (Kurby and Zacks, 2008; Radvansky, 2012; Sargent et al., 2013), such that elements within an event are bound together more cohesively than elements across events (Ezzyat and Davachi, 2011; DuBrow and Davachi, 2013). A natural candidate for mediating the effects of event boundaries on memory is the hippocampus, given multiple findings which, taken together, suggest a general sensitivity to prediction error (e.g., Strange and Dolan, 2001; Köhler et al., 2005; Kumaran and Maguire, 2006; Axmacher et al., 2010; Chen et al., 2011), combined with its well-established role in episodic memory formation (Squire, 1992). Thus during naturalistic experience, event boundaries may be expected to be a particularly strong driver of hippocampal activity (though see Schapiro et al., 2016), potentially registering the preceding event to long-term memory as a bound representation (Ben-Yakov and Dudai, 2011; Richmond and Zacks, 2017).

This hypothesis gains further support from studies that have found increased hippocampal activity at the offset of discrete film clips (Ben-Yakov and Dudai, 2011; Ben-Yakov et al., 2013, 2014) or following a context switch (DuBrow and Davachi, 2016), and have related this activity to subsequent memory. However, the stimuli in these studies were clearly dissociated from one another, with boundaries imposed by the experimenter. It is unknown whether the hippocampus responds to subjective event boundaries during continuous, more naturalistic experience. Baldassano et al. (2017) analysed data from a full-length film and found: i) increased hippocampal responses coinciding with shifts in cortical activity patterns, and ii) a coincidence (35-40% match) of pattern shifts and annotated event boundaries. While these findings hint at a potential link between hippocampal activity and event boundaries, this study did not test a direct link, nor the potentially confounding effects of perceptual change.

Here we examined the direct relationship between hippocampal activity and boundaries, and how that activity is modulated by the proportion of observers indicating a boundary (“boundary strength”), after adjusting for various perceptual confounds. Furthermore, even though hippocampal activity may be <u>sensitive</u> to event boundaries, it may not be <u>specific</u> to boundaries. That is, there may be other time-points with high hippocampal activity that do not correspond to event boundaries. By defining peaks in hippocampal activity during continuous stimuli, we were able to characterise the specificity of the hippocampal response to event boundaries. Finally, we assessed whether sensitivity to boundary strength was <u>selective</u> to the hippocampus by exploring an atlas of other brain regions.

We addressed these questions by analysing functional magnetic resonance imaging (fMRI) data from two independent datasets in which participants watched films (Shafto et al., 2014; Hanke et al., 2016). By combining hypothesis-driven and data-driven approaches, we were able to demonstrate that the hippocampal response is sensitive, selective and specific to event boundaries.

## Materials and Methods

We analysed data from two datasets with fMRI scanning of participants viewing films – Stage II of the Cambridge Centre for Ageing and Neuroscience (*CamCAN*, www.cam-can.org) project (see Shafto et al., 2014 for more details) and the 3T audiovisual movie dataset of the *studyforrest* project (http://studyforrest.org, see Hanke et al., 2016 for more details).

### Participants

#### CamCAN

We used the 253 adults (131 female) who were aged 18-47 (mean age 34.8, SD[standard deviation]= 7.9) from the healthy, population-derived cohort tested in Stage II of the *CamCAN* project (Shafto et al., 2014; Taylor et al., 2017). The majority of participants (228) were definitively right-handed (defined as a handedness measure of >=50 on a scale of −100[Left] to 100[Right]; definitively left-handed were defined as <=-50 and those with a handedness measure of −49-49 were considered undetermined). All participants were native English speakers. Ethical approval was obtained from the Cambridgeshire Research Ethics Committee and all participants gave their written informed consent prior to participation.

#### studyforrest

The current analysis focused on 15 participants (mean age 29.4, range 21–39, 6 female) for whom functional MRI was collected during film viewing (Hanke et al., 2016; Sengupta et al., 2016). The participants were all right-handed native German speakers with normal visual function. Ethical approval was obtained from the Ethics Committee of the Otto-von-Guericke University and all participants gave informed consent prior to participation.

### Experimental Design

#### CamCAN

Participants viewed an abridged version of Alfred Hitchcock’s black-and-white television drama “Bang! You’re Dead” (Hasson et al., 2008, 2010), edited from 30 min to 8 min while maintaining the plot (Shafto et al., 2014). The film was chosen to be compelling but unfamiliar to participants.

#### studyforrest

Participants viewed the film Forrest Gump (R. Zemeckis, Paramount Pictures, 1994) with German dubbing. The film was edited to be 2 hours and divided into 8 segments, each approximately 15 min long, presented in a separate scan (Hanke et al., 2016). All participants except one had previously seen the film (and the additional participant had previously heard an audio-only version).

### Film segmentation

We identified the occurrence of event boundaries using subjective annotations. Five observers viewed each of the films and indicated with a keypress when they felt ‘one event (meaningful unit) ended and another began’ (based on the event segmentation approach in Newtson, 1973; Zacks et al., 2010). In terms of granularity of segmentation (coarse/fine-grained), participants were instructed to segment in the manner that felt most natural to them. To account for response time, 0.9s was subtracted from the logged button presses (calculated based on prior testing to estimate reaction time). Because observers indicated they occasionally pressed the button accidentally, only moments in time identified by at least two observers were defined as event boundaries. If different observers marked boundaries within one time-point (repetition time, TR) of one another, these were treated as a single boundary (and the time of the boundary was defined as the average time identified by the different observers). In *studyforrest*, the TR was shorter and the boundaries more frequent, resulting, following this heuristic, in a minimal temporal distance of 2.02s between boundaries, compared to 4.3s in *CamCAN*. For consistency, an additional iteration was run on the *studyforrest* boundaries, averaging all pairs of boundaries that were less than 4s apart (5 pairs across all runs). In the final set of boundaries used for analysis, the distance between consecutive boundaries in *CamCAN* was 4.3-94.4s (mean 19.3s, SD=18.1s) and 5-140.6s (mean 41.2s, SD=29.4s) in *studyforrest*. The boundaries were recorded using PsychoPy v1.85.0 (Peirce, 2007). In *CamCAN* there were 25 event boundaries (7 were identified by all 5 observers, 7 by 4, 5 by 3 and 6 by 2) and in *studyforrest* there were 167 boundaries (10 were identified by all 5 observers, 24 by 4, 49 by 3 and 84 by 2), 11-28 per run. An advantage of using an independent set of observers for segmentation is that asking participants to indicate boundaries while watching a film in the scanner may alter the brain responses, e.g., by making boundaries task-relevant, and no longer corresponding to naturalistic viewing. An alternative would be to ask each participant to annotate the film again, after having watched it in the scanner, but a concern here, which is of particular relevance for the hippocampus, is that memory for the film might affect the decisions about event boundaries.

### fMRI Acquisition

#### CamCAN

Imaging was performed on a 3T Siemens TIM Trio scanner (Siemens Medical Solutions) at the MRC Cognition and Brain Sciences Unit, employing a 32 channel head coil. High resolution 3D T1-weighted structural images were acquired using a Magnetization Prepared Rapid Acquisition Gradient-Echo (MP-RAGE) pulse sequence (1x1x1 mm resolution, TR[repetition time]=2250ms, TE[echo time]=2.99ms, TI[inversion time]=900ms, flip angle=9°, FOV[field of view]=256x240x192mm, GRAPPA acceleration factor =2). Functional images were acquired using a multi-echo, T2*-weighted echo-planar imaging (EPI) sequence (TR =2470ms, TE[five echoes]=[9.4,21.2,33,45,57ms], flip angle=78°, 32 axial slices with 3.7mm thickness and a 20% gap, FOV=192×192mm, voxel-size=3×3×4.44mm).

#### studyforrest

Imaging was performed on a 3T Philips Achieva scanner (Philips Medical Systems), employing a 32 channel head coil. High resolution T1-weighted structural images (Hanke et al., 2014) were acquired using a 3D turbo field echo (TFE) sequence (acquisition voxel size of 0.7 mm with a 384×384 in-plane reconstruction matrix [0.67 mm isotropic resolution], TR=2500ms, TE=5.7ms, TI=900ms, flip angle=8°, FOV=191.8×256×256mm, bandwidth 144.4 Hz/px, Sense reduction AP 1.2, RL 2.0). Functional images (Hanke et al., 2016) were acquired using a gradient-echo, T2*-weighted echo-planar imaging (EPI) sequence (TR =2000ms, TE=30ms, flip angle=90°, 35 axial slices with 3.0mm thickness and a 10% gap, FOV=240x240mm, voxel-size=3×3×3mm). Slices were automatically positioned in AC-PC orientation using Philips’ ‘SmartExam’ (such that the topmost slice was at the superior edge of the brain).

### Data Preprocessing

Data from both studies were pre-processed using SPM12 (www.fil.ion.ucl.ac.uk/spm), automated with the Automatic Analysis (AA) 4.2 pipeline system (Cusack et al., 2014, for details on the specific analysis used, see Taylor et al., 2017) in MATLAB (v8.5.0 R2015a, The MathWorks). T1 anatomical images were coregistered to the Montreal Neurological Institute (MNI) template using rigid-body transformation, bias-corrected, and segmented by tissue class. Diffeomorphic registration was then applied to the grey matter to create a group template, which was, in turn, affine-transformed to MNI space. For the *CamCAN* functional images, the data from the multiple echos were first averaged, weighted by the contrast-to-noise ratio of each echo for each voxel. For both datasets, the functional images were corrected for motion and then corrected for slice acquisition times by interpolating to the middle slice. The images were rigid-body co-registered to the corresponding T1 image and the spatial transformations from that image to MNI space (diffeomorphic+affine) were applied. Finally, effects of abrupt motion were reduced by applying wavelet-despiking (Patel et al., 2014). In the *CamCAN* data, two additional steps were performed: anatomical segmentation was informed by additional T2 images and field maps were used to correct EPI distortions prior to motion correction (no T2 images or fieldmaps were available for the *studyforrest* data). High-pass filtering (cutoff of 256s) was implemented with a cosine basis set as part of the general linear model (GLM) within SPM12 (for the pattern analyses, the data were filtered first by taking the residuals from a GLM containing just the cosine terms).

### Anatomical Region of Interest (ROI) Definition

The T1 images, after alignment to the group template in MNI space, were averaged separately for each dataset. Bilateral group-ROIs of the hippocampus were then manually traced using ITK-SNAP (www.itksnap.org, Yushkevich et al., 2006). As we found no difference between right/left hippocampus in the effects of interest (and did not have an a priori hypothesis regarding such differences), the analyses were all run on the average across left and right hippocampus. The visual (‘Visual Cortex V1’), auditory (‘Primary auditory cortex TE1.0+TE1.1’) and angular gyrus ROIs were defined using the Juelich atlas (Eickhoff et al., 2005). The angular gyrus ROI was defined using the same ROI as in Baldassano et al. (Baldassano et al., 2017). The whole-brain analysis was run using an ROI approach – running the model on all ROIs from the Harvard Oxford Atlas (Desikan et al., 2006).

### Angular gyrus (AG) pattern shift analysis

In a recent study, Baldassano and colleagues (Baldassano et al., 2017) used a data-driven approach for event segmentation of realistic experience. They found that in high-level regions such as the angular gyrus (AG) and posterior medial cortex, points in time of rapid change in activity patterns (cortical event boundaries) corresponded to event boundaries as defined by human observers. Moreover, cortical event boundaries, particularly in the AG, coincided with an increase in hippocampal univariate activity. Our analysis focused on the AG, using an anatomical ROI from the Eickhoff atlas (Eickhoff et al., 2005). To test whether AG pattern changes could account for any hippocampal response to event boundaries in the current studies, we defined AG boundaries implementing the Hidden Markov Model segmentation from Baldassano et al. (2017), with one alteration. In the original segmentation procedure, the number of events was estimated in a data-driven manner. In the current studies, this approach did not yield optimal results, so we defined the number of events in each scan according to the number of event boundaries. We classified the AG pattern shifts according to whether they matched an event boundary (i.e., occurred up to 2 timepoints after) and separately averaged the hippocampal response around match/non-match AG pattern shifts. The hippocampal response was calculated by z-scoring the entire timecourse for each run of each participant, then averaging over participants, over left/right hippocampus and over event boundaries. An examination of the original results (Baldassano et al., 2017) reveals that the peak hippocampal response occurred in the first 4-5 s after the AG shift. Therefore, we computed the amplitude of the hippocampal response as an average of timepoints 0-2 relative to the AG shift.

## Statistical Analysis

### Assessing significance of response to boundaries

The amplitude of the hippocampal response to event boundaries was measured using a GLM with a single predictor for all event boundaries (i.e, a stick function at event boundaries, convolved with the canonical hemodynamic response function [HRF]), together with the high-pass filter regressors as nuisance predictors. To assess the significance of the hippocampal response to boundaries, we compared it with the measured response to boundaries in 1000 random permutations of the event durations (the intervals between consecutive boundaries), and used the ratio of permutations with a larger response than the intact one (in absolute value) to derive a p-value (Figure 1A). A similar approach was used to assess the significance of the hippocampal response at AG pattern shifts, comparing the amplitude of the response to the amplitude calculated when permuting events (here defined as the epochs between AG pattern shifts).

**Figure 1.**
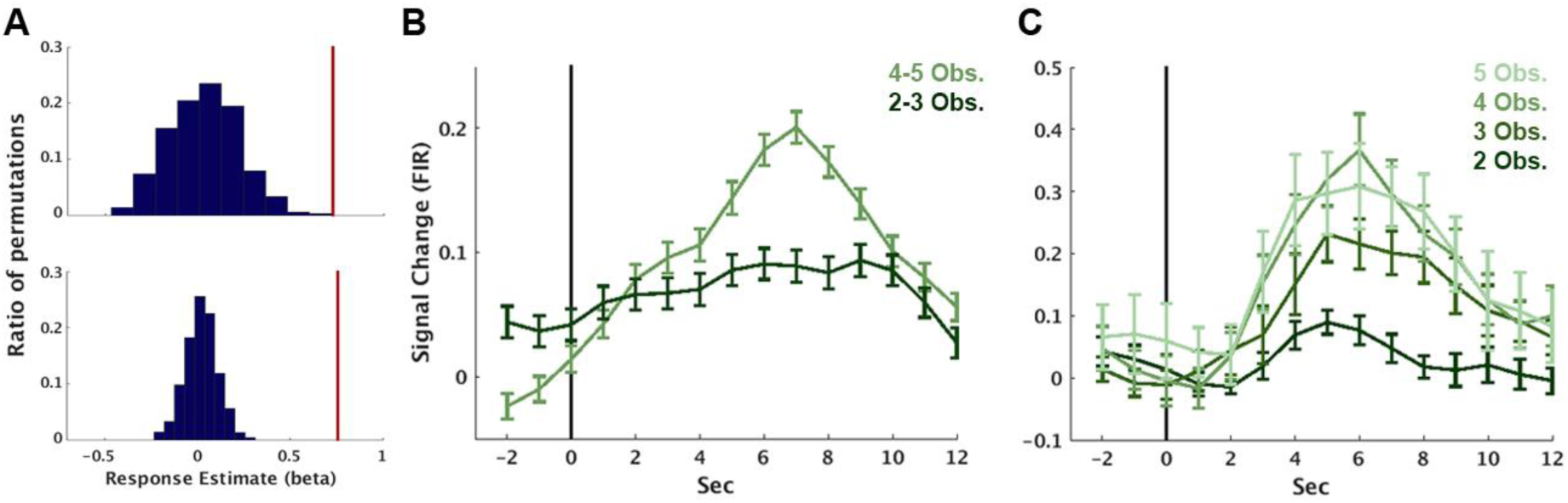
Hippocampal response to event boundaries. (A) The average amplitude of the canonical response to an event boundary (red lines) relative to the distribution of responses when randomly shuffling event order. Shown for *CamCAN* (top) and *studyforrest* (bottom). (B,C) Average response, across participants, to event boundaries in *CamCAN* (B) and *studyforrest* (C). The per-participant time-course was calculated using an FIR, and error bars reflect the standard error of the mean at each timepoint. The vertical black line represents the event boundary.

### Mixed-effects model

When using films as memoranda, the *stimulus-as-fixed-effects fallacy* (Clark, 1973; Westfall et al., 2016) becomes more pertinent, as each film has specific characteristics that do no generalise to all films. We thus used a mixed-model for statistical analysis, incorporating both participants and items (event boundaries) as random effects (Baayen et al., 2008). We first estimated the hippocampal response to each participant-by-item in a GLM with a separate predictor for each boundary of each participant, as well as high-pass filter confound predictors. All items closer than 6s to one another or less than 10s from the end of the run were removed from subsequent analysis, leaving 22/25 event boundariess in *CamCAN* and 160/167 in *studyforrest*. The resulting betas were then submitted to a linear mixed-model with nObservers as the (fixed) effect of interest, indicating the number of observers who identified the item as an event boundary, and participant and item as random effects. Additional fixed effects included handedness in *CamCAN* (Right/Left/Undetermined) and run number in *studyforrest* (for each of the 8 runs).

The linear models were then fitted using restricted maximum likelihood with the lme4 (Bates et al., 2014) and lmerTest (Kuznetsova et al., 2017) packages in R (R 3.1.3, R Core Team, 2017, https://www.R-project.org/),> with the following formulas (‘*’ indicates interaction, and ‘(1|x)’ indicates a random effect):

<u>CamCAN</u>: ‘betas ∼ nObservers + handedness + nObservers*handedness + (1|participant) + (1|event boundary)’

<u>studyforrest</u>: ‘betas ∼ nObservers + runNum + (1|participant) + (1|event boundary)’

Because event boundaries are typically characterised by various types of visual and auditory change, we ran a second analysis incorporating multiple confound predictors estimating the following attributes:

<u>isLoc/isTemp (change in location/time)</u> – Event boundaries are often characterised by a change in location, time or both. Due to the sensitivity of the hippocampus to both time and space, we added predictors to account for these changes. For the *CamCAN* film, location/temporal changes were identified by the authors and incorporated into a single predictor, isLocTemp (as the two always coincided). The *studyforrest* project already includes annotations for every film shot, including the spatial location and indication of temporal progression relative to the preceding shot (Häusler and Hanke, 2016). These were used to create separate predictors for change in location (isLoc) or time (isTemp).

<u>visDist (visual distance)</u> – The visual distance between each pair of frames (up to 1500 frames/1 min apart) was calculated using IMage Euclidean distance (IMED, Liwei Wang et al., 2005). This measure takes into account the similarity not only with the parallel pixel in the second image, but also similarity with surrounding pixels (weighted by a distance function), and is thus less sensitive to small movements between frames. We first resized the images to 1/8 of the original resolution (resulting in 96x72 pixels for *CamCAN* frames and 192x82 pixels for *studyforrest* frames), and then compared each pair of frames with the same parameters (distance weighting matrix G and width parameter σ) used in Wang et al. (Liwei Wang et al., 2005), with one exception: For ease of calculation, only pixels in a 9X9 square around a given pixel were taken into consideration, as beyond this range the weights in the G matrix were virtually zero. The original distance measure is for grayscale images, summing the distance of all pixels to calculate the global image distance. To extend the measure for colour images (in *studyforrest*), we calculated the distance for each channel of each pixel separately, then summed over channels and pixels for the global measure.

After having calculated the visual distance between pairs of frames, the distance across each boundary was defined as the maximal distance between any frame in the 1s window before the boundary with any frame in the 1s window following the boundary. The same approach to calculation visual change across boundaries was applied to all following visual measures.

<u>visCorr (visual correlation)</u> – The visual correlation between each pair of frames (up to 1500 frames/1 min apart) was calculated using IMage Normalized Cross-Correlation (IMNCC, Nakhmani and Tannenbaum, 2013). This measure uses a similar approach to IMED for calculating correlations while taking into account spatial relationships of pixels. When calculating IMNCC we used the same parameters (G, σ) as for the IMED calculation above.

<u>visHistDist (visual histogram distance)</u> – In addition to measuring the visual distance between frames, we measured the visual distance between the histograms of the frames. The rationale for this is that two frames may be quite distinct in terms of the objects they contain and their spatial layout, but still depict a similar setting, with similar lighting and colours. To test for such global similarity, we calculated the histogram of each frames and computed the Euclidean distance between the histograms. For RGB frames (*studyforrest*) we calculated the histogram for each channel separately, then computed the distance over all bins of the three histograms together.

<u>lumDist (luminance distance)</u> – In order to detect global lighting changes, we calculated the difference in global luminance between frames – first calculating the average luminance over all pixels then taking the absolute difference.

<u>DCNN (deep convolutional neural network)</u> – DCNNs are able to extract higher-order information from images, beyond the low-level perceptual properties, with higher layers corresponding to higher-order visual regions (Güçlü and van Gerven, 2015). To automatically identify similarities between frames at multiple levels of visual feature hierarchy, we submitted each frame to AlexNet, one of the most commonly used DCNNs for image identification (Krizhevsky et al., 2012). We then correlated the representation of each pair of frames in each layer of the network. Calculation of the correlation across boundaries was identical to the rest of the visual features, with the exception that every boundary had 21 correlation values, one for each layer of the network, yielding 21 vectors of per-boundary correlations. As these vectors were highly correlated, we ran singular value decomposition on the correlation matrix and used the set of first components that explained 90% of the variance in each study (6 in *CamCAN* and 7 in *studyforrest*).

<u>psdCorr (power spectral density correlation)</u> – To assess the auditory difference across boundaries, we calculated the power spectral density (PSD) in the 500ms epochs before and after each boundary, with a cutoff frequency of 5000Hz (when examining the entire audio, over 99% of the power was below this cutoff). The pre-and post-boundary PSDs were correlated as a measure of auditory similarity.

<u>psdDist (power spectral density distance)</u> – In addition to measuring the audio correlations, we calculated the Euclidean distance between the PSDs. absVolDiff (absolute volume distance) – To detect abrupt volume changes we calculated the average volume in the 100ms before and after the boundary, taking the absolute difference as the measure of volume change.

<u>V1Betas, A1Betas</u>: To account for additional low-level visual/auditory changes that may not be captured by the stimulus-defined predictors, we added the per-trial activity in V1 and A1. The per-trial activity was estimated over all participants, in order to extract the stimulus-driven component of their activity.

<u>isAG (angular gyrus boundary)</u> – To account for the effect of AG pattern shifts, we classified the event boundaries according to whether they matched an AG pattern shift (occurred up to 2 timepoints before it) and added this as a binary predictor.

The models with confounds were then fitted using the following formulas:

<u>*CamCAN*</u>: ‘betas ∼ nObservers + handedness + nObservers*handedness + isLocTemp + visDist + visCorr + visHistDist + lumDiff + DCNN[1…6] + psdCorr + psdDist + absVolDiff + V1Betas + A1Betas + isAG + (1|participant) + (1|event boundary)’

<u>*studyforrest*</u>: ‘betas ∼ nObservers + runNum + isLoc + isTemp + visDist + visCorr + visHistDist + lumDiff + DCNN[1…7] + psdCorr + psdDist + absVolDiff + V1_betas + A1_betas + isAG + (1|participant) + (1|event boundary)’

Significance of the predictor of interest (nObservers) was estimated using the anova function of the lmerTest package (Kuznetsova et al., 2017), with type III error calculation and the Satterthwaite approximation for degrees of freedom (Satterthwaite, 1941), found to yield optimal p-value estimations for mixed models (Luke, 2017). The effect size (marginal R^2^) was calculated using the r.squaredGLMM function of the MuMIn package (Barton 2016, MuMIn: Multi-Model Inference. R package version 1.15.6, https://CRAN.R-project.org/package=MuMIn; see Nakagawa and Schielzeth, 2013 for a discussion of this approach).

## Plotting

### Finite Impulse Response (FIR) analysis

The time-course of the average response to each condition was calculated using a FIR analysis of the hippocampal ROI. We extracted and normalised (z-score) the time-course from the hippocampal ROI, and then interpolated the time-course to be in 1s resolution for both projects. Event boundaries were binned according to the number of observers who identified them (2-5 observers). As there were few event boundaries in *CamCAN*, the division was only to high/low (4-5/2-3 observers). We constructed a GLM with a separate predictor for each condition X timepoint in the range [-2…12s] relative to stimulus onset at time 0. This yielded an estimate of the per-condition response for each participant, which was used for plotting purposes (Figure1B,C). The FIR analysis is participant-based, and only used for demonstration purposes (with the mixed analysis used for statistical purposes).

### Model fit plotting

To demonstrate the effect of nObservers on the hippocampal response (Figure 2), we constructed a mixed model consisting of nObservers as a categorical variable, and with participants and items as random effects. We then calculated the response magnitude (model fit) and standard error using the lsmeansLT function of the lmerTest package (Kuznetsova et al., 2017) in R (R 3.1.3, R Core Team, 2017, https://www.R-project.org/) in order to obtain a standard error estimate for plotting that incorporates the variance across both participants and items.

**Figure 2.**
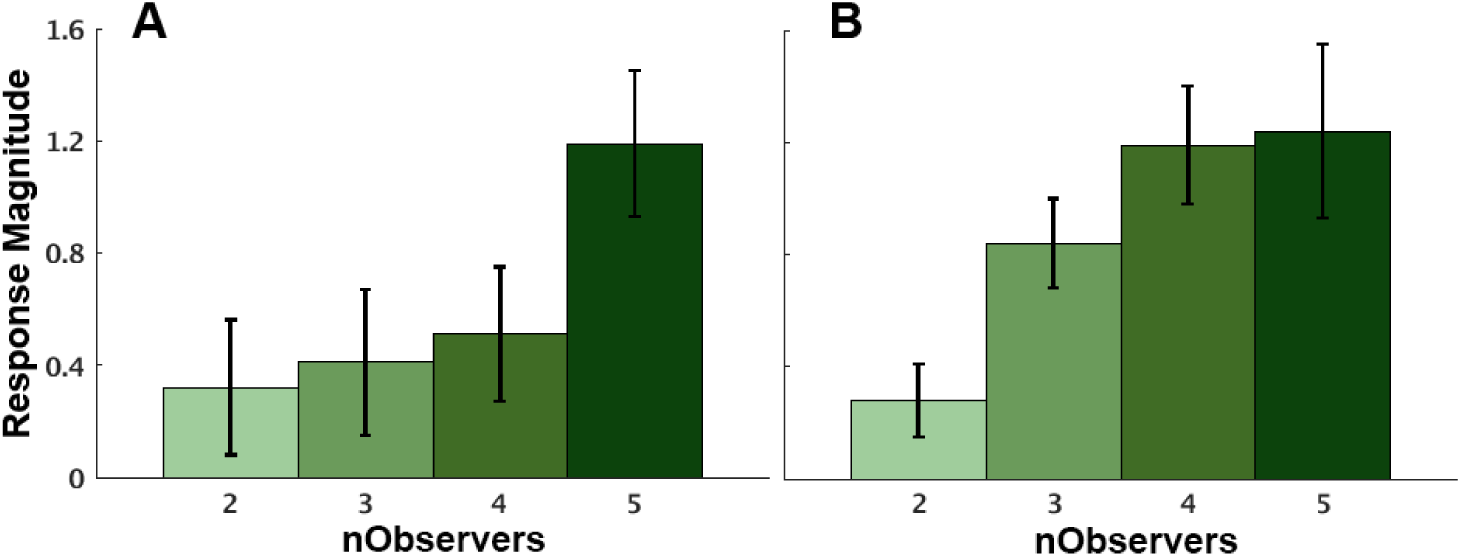
Sensitivity of hippocampal response to boundary strength. The magnitude of the canonical response to event boundaries, binned according to the number of observers that marked them (boundary strength). The response magnitude was calculated as the model fit when modelling number of observers who marked as a boundary (nObservers) separately (equal to the average over participants and boundaries) and errorbars depict the standard error of the fit (incorporating variance across participants and event boundaries). Results are presented for *CamCAN* (A) and *studyforrest* (B).

### Hippocampal data-driven segmentation

In addition to the hypothesis-driven approach, we defined “hippocampal-events” as the set of momentary events that, when modelled, minimised the residual error in the hippocampal time-course. The raw hippocampal time-course of each participant (averaged over all voxels in the hippocampus) was first high-pass filtered and z-scored, then averaged across participants. The number of hippocampal-events was pre-defined as the number of event boundaries, and we iteratively chose hippocampal-events until this number was reached (in *studyforrest* the number of event boundaries was calculated per-run). Hippocampal-events were chosen in the following manner, starting with a general linear model, M, that included only a constant predictor and an empty list of hippocampalevents:

1) Run over all timepoints that have not yet been added to the model.

2) For each timepoint – create a temporary model M_temp_ by adding a predictor with a stick function at that timepoint, convolved with the canonical HRF, and calculate the residual sum of squares (RSS) of this model.

3) Choose the timepoint (model) that most reduces the RSS, and define it as a hippocampal-event, adding its corresponding predictor to M before the next iteration. This was done under two constraints – the beta value associated with the hippocampal-event is positive, and does not flip the signs of any beta values of the hippocampal-events defined in previous iterations.

To calculate the significance of the match between the event boundaries (hypothesis-driven) and hippocampal-events (data driven), we created two binary vectors: 1) a vector with 1 at the timepoints of event boundaries and 2) a vector with 1 at hippocampal-events and each timepoint preceding/following it. The phi coefficient (Pearson correlation for binary variables) between these vectors was taken as the measure of contingency between these two vectors, such that events separated by up to one timepoint were considered a match.

## Results

Our main question of interest was whether subjective event boundaries trigger hippocampal activity during continuous naturalistic experience. This entails estimating both the sensitivity and specificity of hippocampal activity to event boundaries. Sensitivity was assessed in a hypothesis-driven approach, by examining the hippocampal response at event boundaries subjectively-annotated by independent observers. We also assessed the selectivity of the hippocampal response by comparing hippocampal sensitivity to event boundaries with that of other anatomical regions across the whole brain. Specificity was assessed using a data-driven approach, identifying hippocampal-events based on the amplitude of the hippocampal response and testing the overlap between these events and subjective boundaries.

### The hippocampus is sensitive to event boundaries

In the hypothesis-driven analysis, we defined event boundaries by using a separate group of 5 people who indicated with a keypress when they experienced an event shift while watching each film (see Methods). We first assessed overall sensitivity to the occurrence of event boundaries by comparing the estimated response to a boundary to the distribution of estimated responses when shuffling the events (the durations between boundaries). In both studies, the estimated response to boundaries in the intact event order was larger than for any of the random permutations (equivalent to p<0.001, Figure 1A). Having determined overall sensitivity to boundaries, we set out to determine whether the hippocampal response was modulated by the number of observers who marked the boundary, as a measure of “boundary strength”. For demonstration purposes, we first plotted the time-course of the average response to a boundary, binned according to the number of observers who marked it (ranging from 2-5 observers, though in *CamCAN*, boundaries were only binned into two levels due to the small number of event boundaries). In both studies we found a higher response when a larger number of observers marked a moment in time as a boundary (Figure 1B,C). To quantify this effect, we ran mixed-effects models with nObservers as the (fixed) effect of interest, participant and event boundaries as random effects, as reported below for each film separately.

### CamCAN

We found a monotonic increase in the hippocampal response to a boundary as a function of the number of observers who marked that boundary (Figure 2A), and the mixed-effects model revealed this modulation was significant (p=0.01, F(1,50.5)=7.2). The effect size was small (R^2^=0.02), but it is worth noting the model was run on single trial estimates, and thus effect sizes are expected to be small compared to models which average over participants or within-condition items.

Because event boundaries are typically characterised by visual and auditory shifts, we fit an additional model to test whether low-level visual and auditory changes could account for the hippocampal sensitivity to boundaries. These included various measures of visual and auditory change across the boundary, such as luminance and sound-level differences, as well as responses extracted from early visual and auditory cortices (for a full list, see Methods). In addition, we added confound predictors to account for two alternate hypotheses regarding the trigger for hippocampal responses. First, a recent study (Baldassano et al., 2017) found increased hippocampal activity following boundaries defined by cortical pattern shifts, particularly in the angular gyrus (AG). These cortical pattern changes exhibited a large degree of overlap with subjectively-annotated boundaries, thus hippocampal sensitivity to event boundaries could arise from AG pattern shifts. To test this, we added to the model a binary predictor indicating whether an AG boundary occurred in temporal proximity to each annotated one. Second, we added a predictor of change in time/space (isLocTemp) to determine whether the sensitivity to boundaries was driven by such changes. When adding all confounds, the effect of nObservers was no longer significant (p=0.74, F(1,3.4)=0.13), potentially due to the large number of confounds (17) relative to the number of event boundaries (22). Indeed, when adding each confound predictor to the model separately, the effect of nObservers remained significant, except when adding isLocTemp (for one of the predictors, estimating difference in luminance, there was only a trend, with p=0.07). Notably, most event boundaries were characterised by a change in location such that the predictors of isLocTemp and nObservers were highly dependent (ϕ=0.65), and their respective contributions could not be properly dissociated. Thus, we cannot conclude, based on this dataset alone, whether this modulation by boundary strength may be accounted for by other perceptual factors, particularly spatial/temporal change.

#### Studyforrest

In *studyforrest* we similarly found a monotonic increase with the number of observers (Figure 2C, p=6.9x10^−7^, F(1,151)=26.9, R^2^=0.02). Even when adding all confounds, including separate confounds for spatial and temporal changes (isLoc, isTemp), the effect remained significant (p=0.004, F(1,132)=8.6). Notably, the hippocampus also demonstrated sensitivity to change in location (p=0.04, F(1,134)=4.5, when accounting for all confounds except time) or time (p=0.006, F(1,134)=7.7, when accounting for all confounds except location), but not when accounting for nObservers (p=0.17, F(1,133)=1.9 for location change; p=0.1, F(1,133)=2.7 for temporal change). Thus, while changes in location/time modulate hippocampal activity, this does not seem to account for its sensitivity to boundary strength.

In both studies, we found that the hippocampus is sensitive to boundary strength (the number of observers who subjectively reported a shift). Furthermore, in *studyforrest*, where there was a sufficient number of event boundaries to assess the relative contribution of potential confounds, we found that the sensitivity to the number of observers could not be explained by objective measures such as visual/auditory change or by change in location/time. The modulation of hippocampal response by boundary strength suggests the hippocampus is not only sensitive to the occurrence of event boundaries, but also to their salience. However, since we do not have event boundary annotations from the fMRI participants, we cannot distinguish between two possibilities: i) that the hippocampus responds in a graded manner depending on boundary strength, or ii) that it responds in a binary manner within each participant, and that more salient boundaries occurred for a larger proportion of the scanned participants. Nonetheless, there are good reasons why one might be wary of using boundaries indicated by individual participants for this purpose (see Methods).

#### Both hippocampal activity and AG patterns are driven by event boundaries

Another potential explanation for the increase in hippocampal activity at event boundaries is that event boundaries elicit cortical pattern shifts, which in turn drive hippocampal activity (Baldassano et al., 2017). While the addition of the AG predictor did not account for the sensitivity to boundary strength in the above analyses, AG pattern shifts could still account for the overall hippocampal response to boundaries, regardless of their boundary strength. To test this, we divided AG pattern shifts into those that corresponded with an event boundary (AG-match) and those that did not (AG-non-match), averaging the hippocampal response around each type (Figure 3). We replicated the finding of Baldassano and colleagues (Baldassano et al., 2017) of an increased hippocampal response to overall AG pattern shifts (*CamCAN*: p=0.01, *studyforrest*: p=0.002). However, this increase was only found for AG pattern shifts that coincided with event boundaries: For matching shifts (13/25 in *CamCAN* and 38/167 in *studyforrest*), there was a significant increase in hippocampal activity at the shift (*CamCAN*: p=0.002, *studyforrest*: p<0.001, higher than all random permutations), whereas for the non-matching shifts there was no significant increase in hippocampal activity (*CamCAN*: p=0.63, *studyforrest*: p=0.42). In a direct comparison of match and non-match, the difference was significant (*CamCAN*: p=0.01, *studyforrest*: p<0.001) in both studies. This suggests that both AG pattern shifts and increased hippocampal activity are driven by the occurrence of event boundaries, rather than AG pattern changes and hippocampal activity being directly related.

**Figure 3.**
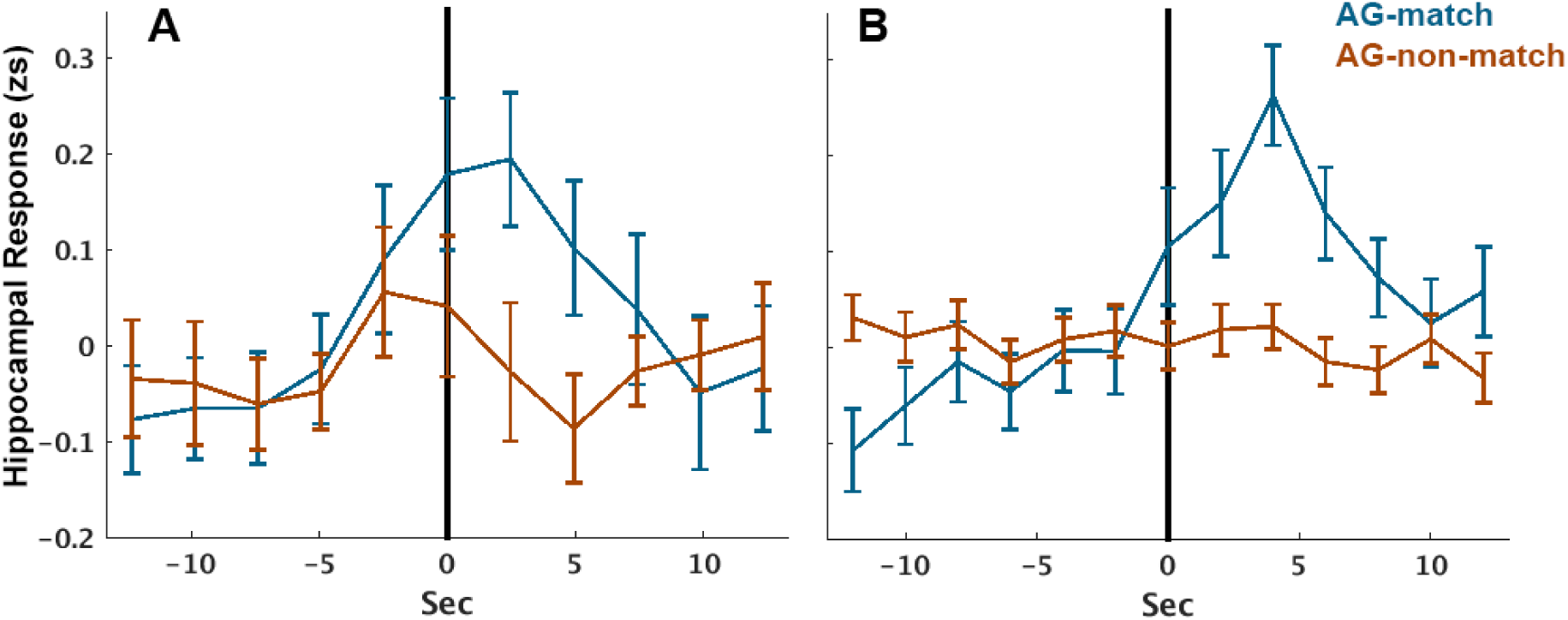
Hippocampal response to AG pattern shifts. The average zscored (zs) hippocampal response at AG pattern shifts that match/do not match annotated event boundaries. Time zero (vertical lines) represent the time of the pattern shift, uncorrected for hemodynamic delay. Error bars represent standard error of the mean (across pattern shifts). Results are shown for *CamCAN* (A) and *studyforrest* (B).

### Selectivity of hippocampal modulation by boundary strength

Our a-priori interest, based on previous studies, was the effect of event boundaries on hippocampal activity. However, additional regions may show similar modulation by nObservers, given the large number of regions that have been reported to respond to event boundaries in general (Zacks et al., 2001, 2010). We tested this by re-running the above mixed-model effects on anatomically-defined regions across the brain taken from the Harvard-Oxford Atlas (Desikan et al., 2006). We averaged across left and right homologous regions. Of the 55 homologous regions in the atlas, five showed a significant modulation by nObservers in both experiments, with the hippocampus exhibiting the most significant modulation of these five regions in both experiments. Moreover, the hippocampus was the only region to show a monotonic increase in the two experiments. Thus while some regions were modulated by nObservers in one experiment or the other, the only region that appeared to exhibit a consistent and significant monotonic modulation in both experiments was the hippocampus, demonstrating that it is relatively selective in its response to boundary strength. The selectivity of the hippocampus suggests that its modulation by nObservers is more likely to reflect sensitivity to the strength of the boundary (rather than a binary response aggregated over a larger number of participants), as multiple regions that have previously been found to respond to boundaries (Zacks et al., 2001, 2010), do not show this increase with nObservers.

### Specificity of hippocampal response to event boundaries

In the data-driven analysis, we set out to reveal whether increased hippocampal activity is specific to event boundaries. To do so, we defined “hippocampal-events” as points in time that, when modelled as events, best explained the observed hippocampal response, with the number of hippocampal-events set according to the number of event boundaries. These hippocampal-events were then compared to the pre-defined boundaries, dividing them into those that matched a pre-defined boundary (temporal distance of up to 1 TR) versus those that did not (Figure 4). In *CamCAN*, 15/25 hippocampal-events matched pre-defined boundaries (60%, ϕ=0.19, p=0.01). Of these, 9 matched strong boundaries (4/5 observers, 64% of such boundaries) and 6 matched weak boundaries (2/3 observers, 54%). In *studyforrest*, 54/167 hippocampal-events matched pre-defined boundaries (32%, ϕ=0.13, p=4.9x10^−14^). Of these, 20 matched strong boundaries (59% of such boundaries) and 34 matched weak ones (26%).

**Figure 4.**
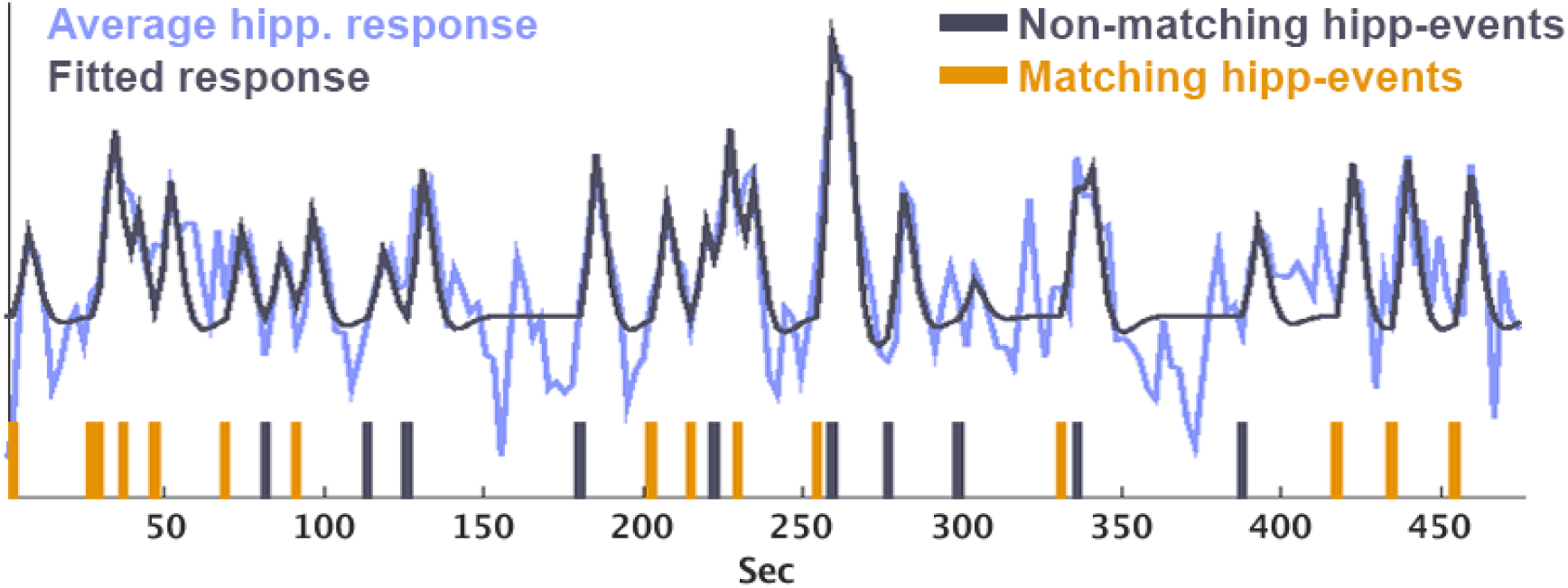
Specificity of hippocampal-events (data-driven) to predefined event boundaries. The average hippocampal time-course in *CamCAN* plotted together with the fitted model. The vertical lines indicate the hippocampal-events estimated from the data – the set of events that minimised the residual error of the model when fitting to the hippocampal time-course. The model was created by convolving each of these events with an HRF as a separate predictor, yielding the fitted model plotted. The hippocampal-events were then divided into those matching a pre-defined boundary (up to 1 TR from a boundary, in orange, 60% of hippocampal-events) and non-matching ones (grey).

Taken together, these complementary analyses suggest the hippocampus is both sensitive and specific in its response to the occurrence of event boundaries. Moreover, its response is monotonically modulated by boundary strength, a pattern selective to the hippocampus.

## Discussion

We examined the relationship between event boundaries in continuous experience and the brain’s response measured by fMRI, using films as a proxy for real-life experience. In particular, we examined the sensitivity, specificity and selectivity of the response of the hippocampus, given extant but indirect evidence implicating it in processing of event boundaries (Ben-Yakov and Dudai, 2011; Ben-Yakov et al., 2013; DuBrow and Davachi, 2016; Baldassano et al., 2017). In two distinct films, subjective event boundaries were defined by an independent group of observers.

Event boundaries were a reliable trigger for an increased hippocampal response. Moreover, the hippocampal response was sensitive to boundary strength, with the strongest hippocampal response occurring at boundaries identified by all five observers. Interestingly, in the longer film (*studyforrest*, which had a sufficient number of events), this sensitivity remained after covarying a large number of measures of perceptual change at those event boundaries. To address selectivity, we examined the responses of a large number of other brain regions, and found that the hippocampus was the only region that showed significant monotonic sensitivity to boundary strength for both films. Finally, to address specificity, we took an alternative, data-driven approach, in which we identified moments in which the hippocampus exhibited the strongest responses, and tested the correspondence between these “hippocampal-events” and the subjective event boundaries. In both films, there was a significant match, reaching 60% in *CamCAN*.

Our finding that the hippocampus is sensitive to subjective event boundaries complements other studies that explicitly manipulated boundaries using discrete stimuli, such as film clips (Ben-Yakov and Dudai, 2011; Ben-Yakov et al., 2014) or sequences of pictures (DuBrow and Davachi, 2014, 2016; Hsieh et al., 2014) and studies identifying hippocampal sensitivity to spatial boundaries (e.g., Doeller et al., 2008; Bird et al., 2010; Gupta et al., 2012; McKenzie and Buzsáki, 2016). It is difficult however to determine how much of the hippocampal response in these experiments relates to perceptual change at discrete points in time, rather than subjective segmentation of continuous stimulation. Milivojevic et al. (2016) examined hippocampal activity in a continuous film, but focused on the representation of the events themselves, rather than sensitivity to boundaries. The only prior study, to our knowledge, that examined hippocampal activity to subjective event boundaries during continuous films is that by Baldassano et al (2017). These authors found an increase in hippocampal activity that coincided with shifts in cortical patterns (e.g, in angular gyrus). These cortical pattern shifts also tended to coincide with subjective boundaries, revealing an indirect link between hippocampal activity and event boundaries in naturalistic experience. Here we provide more direct evidence that hippocampal activity is driven by subjective event boundaries. Indeed, our data do not support the alternative interpretation that the hippocampal response is driven by pattern shifts in cortical regions, because we found an increase in hippocampal activity only at those cortical pattern shifts that coincided with an annotated boundary, suggesting it is the boundaries, and not the pattern shifts, that drive hippocampal activity.

### Caveats

The hippocampal sensitivity to boundary strength did not appear to reflect purely the degree of perceptual change within the film, given that we covaried out a large number of measures of visual and auditory change, including responses in early sensory cortices. Adding these as covariates, as well as explicit changes in location or time, did remove the significant effect of boundary strength in the *CamCAN* film, but not in the *studyforrest* film. Note that we are not claiming that perceptual changes do not contribute to hippocampal responses; only that they are insufficient to account for the full range of hippocampal response to subjective boundaries: The hippocampus responded at moments in time not characterised by a large perceptual change, while salient perceptual changes went ‘unnoticed’ by the hippocampus if they weren’t experienced as a boundary. We discuss below which feature of event boundaries, other than perceptual change, may constitute the primary driver of the hippocampal response.

While the hippocampus was the only brain region (of those in the atlas we used) that showed a significant monotonic relationship with boundary strength in both experiments, we are not claiming that other brains are insensitive to event boundaries. Indeed, many brain regions respond to the occurrence of event boundaries (Zacks et al., 2001, 2010), but they did not show convincing evidence for additional sensitivity to the strength of those boundaries in the present study.

Finally, while the specificity of the data-driven hippocampal-events to subjective boundaries was highly significant, the absolute match was relatively low for the *studyforrest* film (with approximately 30% of hippocampal-events corresponding to subjective boundaries). This lower correspondence relative to the CamCAN film could owe to i) the lower number of participants, rendering peaks in the average hippocampal response more prone to random noise, and/or ii) a difference in the nature of the films (the boundaries in *studyforrest* tended to be less clear-cut due to the narration). An examination of periods of the film around the occurrence of hippocampal-events that did not coincide with boundaries did not reveal a clear trigger, and investigation of a wider array of films may identify additional triggers.

### Functional significance

Two main questions arise as to the nature of hippocampal sensitivity to event boundaries: what constitutes a boundary for the hippocampus, and what type of hippocampal processing does the boundary-triggered activity reflect? With regard to what defines a boundary, Event Segmentation Theory (Zacks et al., 2007) postulates that boundaries correspond to spikes in prediction error. However, in naturalistic experience, these spikes are typically associated with both increased change and greater uncertainty, and it is difficult to disentangle these two (Richmond and Zacks, 2017). Moreover, prediction error can occur for different features of the event (Zwaan et al., 1995; Huff et al., 2014), such as location, time, action, etc, which may have additive effects on the probability of event segmentation (Zwaan et al., 1995; Zwaan and Radvansky, 1998; Magliano et al., 2001; Zacks et al., 2009; Huff et al., 2014). Magliano et al. (2001) found that changes in time or movement, but not location, were sufficient to induce an event boundary, and Magliano et al. (2011) proposed that action discontinuity was the primary driver of segmentation. The types of change that induce segmentation may depend on the nature of the stimulus, and in films specifically, may depend on the types of continuity editing applied (Magliano and Zacks, 2011; Baker and Levin, 2015). Indeed, through bespoke editing rules designed to create a sense of continuity, large changes can go unnoticed (Smith and Henderson, 2008; Smith et al., 2012; Baker and Levin, 2015). Thus, perhaps hippocampal boundaries are elicited not by the degree of perceptual discontinuity, but by the sense of conceptual discontinuity that they elicit. For example, a character joining/leaving a conversation may constitute a boundary, eliciting a hippocampal response despite little perceptual change, whereas a cut to a visually-distinct frame may elicit no response if it is experienced as part of the same event.

The second question pertains to the functional significance of the hippocampal boundary response. Multiple studies have demonstrated that occurrence of boundaries during encoding shapes the subsequent organisation of information in long-term memory (Kurby and Zacks, 2008; Radvansky, 2012; Sargent et al., 2013). For example, episodic elements occurring within an event are bound together more strongly than those encountered across events (Ezzyat and Davachi, 2011; DuBrow and Davachi, 2013). While we did not have measures of memory performance in the current study, an intriguing possibility is that hippocampal activity *during* the event represents the content of that event (Hsieh et al., 2014; Allen et al., 2016; Milivojevic et al., 2016; Terada et al., 2017), and the increased amplitude of that activity at an event boundary reflects registration to long-term memory of a bound representation of the preceding event (Ben-Yakov and Dudai, 2011; Richmond and Zacks, 2017). This is supported by previous evidence that the hippocampus responds more strongly at the offset of (but not during) subsequently remembered versus subsequently forgotten film clips (Ben-Yakov and Dudai, 2011), combined with the role of the hippocampus in episodic binding (Staresina and Davachi, 2009). Notably, a parallel is found in retrieval: the hippocampus is involved in retrieval across event boundaries but not in within-event retrieval (Swallow et al., 2011). Further research will be required to elucidate the exact nature of the hippocampal response, which may signal the context shift itself, thereby leading to segmentation in long-term memory (Polyn et al., 2009; Dubrow et al., 2017), or drive rapid replay of the preceding event, creating a cohesive representation (Ben-Yakov and Dudai, 2011; Sols et al., 2017).

In summary, there has been growing interest the neural basis of memory for naturalistic experience. While less controlled than typical laboratory studies (e.g., in terms of timing), continuous stimuli are closer to real-life memory. Here we demonstrate that the hippocampus is sensitive, specific and selective to the occurrence of event boundaries while watching films. Thus the hippocampus appears important for segmenting continuous experience, most likely in order to transform continuous experience into representations of discrete events for registration into memory.

## Acknowledgements

This work was supported by the UK Medical Research Council (SUAG/010 RG91365) and a Marie Curie Individual Fellowship (705108) awarded to ABY. The research would not have been possible without the data provided by the Cambridge Centre for Ageing and Neuroscience (Cam-CAN) and the studyforrest project. We thank Michael Hanke for his help with the studyforrest dataset, Christopher Baldassano for support with the cortical pattern shift analysis, and Katherine Storrs for her help with the convoluted neural net analysis. We also thank Roni Tibon, Andrea Greve, Alex Kaula and Alex Quent for valuable feedback.

